# Hair follicle stem cell progeny heal blisters while pausing skin development

**DOI:** 10.1101/2020.04.08.032326

**Authors:** Yu Fujimura, Mika Watanabe, Kota Ohno, Yasuaki Kobayashi, Shota Takashima, Hideki Nakamura, Hideyuki Kosumi, Yunan Wang, Yosuke Mai, Andrea Lauria, Valentina Proserpio, Hideyuki Ujiie, Hiroaki Iwata, Wataru Nishie, Masaharu Nagayama, Salvatore Oliviero, Giacomo Donati, Hiroshi Shimizu, Ken Natsuga

## Abstract

Injury in adult tissue generally reactivates developmental programs to foster regeneration, but it is not known whether this paradigm applies to growing tissue. Here, by employing blisters, we show that epidermal wounds heal at the expense of skin development. The regenerated epidermis suppresses the expression of tissue morphogenesis genes accompanied by delayed hair follicle (HF) growth. Lineage tracing experiments, cell proliferation dynamics, and mathematical modeling reveal that the progeny of HF junctional zone stem cells, which undergo a morphological transformation, repair the blisters while not promoting HF development. In contrast, the contribution of interfollicular stem cell progeny to blister healing is small. These findings demonstrate that tissue development can be sacrificed for the sake of wound regeneration and suggest that tissue repair does not coincide with the reactivation of developmental programs in all regenerative contexts. Our study elucidates the key cellular mechanism of wound healing in skin blistering diseases.

## Introduction

Tissue responds to injury by transforming its cellular components and extracellular matrix from homeostasis into a regenerative state. Damaged tissue typically reactivates an embryonic gene program in epithelia to accelerate tissue regeneration (1-4). However, it is unknown whether this phenomenon also applies to injuries in developing tissue, in which the embryonic gene expression program is switched on before damage.

The epidermis is a stratified epithelium of the skin and is located on the surface of the body, where it serves as a barrier against external stimuli and microorganisms (5). Cellular proliferation and differentiation in the epidermal basal layer, where epidermal stem cells (SCs) are present, are fine-tuned to maintain the integrity of the epidermis (6). The epidermis attaches to the dermis through proteins in the epidermal basement membrane zone (BMZ) (7). Epidermal BMZ proteins function as a niche for epidermal SCs (8), and the loss of these proteins, such as α6 integrin (ITGA6), β1 integrin, and collagen XVII (COL17), leads to transient epidermal proliferation (9-11).

Skin wounding causes pain and carries a significant risk of bacterial infection. The sources of skin wound healing have been extensively investigated in experimental animals (12, 13). Hair follicles (HFs), epidermal appendages, fibroblasts, and immune cells coordinate to heal the wound, and the contribution of each component can vary depending on the assay (14). Wounding in adult skin induces the expression of genes regulating epidermal development, including SOX11 and SOX4 (1).

Conventional skin wounding assays have employed full-thickness skin wounds, in which all skin components are removed, including the epidermis, epidermal appendages, dermis, and subcutaneous fat tissue. In contrast to conventional full-thickness skin wounds, epidermal detachment, as exemplified by subepidermal blisters, is distinctive because it does not affect the structures below the epidermis per se. The epidermis is detached from the dermis in several pathological conditions, such as burns (15), congenital defects in epidermal BMZ proteins (epidermolysis bullosa (EB)) (16, 17), autoimmunity to these proteins (pemphigoid diseases) (18), and severe drug reactions, such as Stevens-Johnson syndrome/toxic epidermal necrolysis (19). Although the cells that contribute to the repair of full-thickness skin wounds have been identified (13, 20-24), the cellular dynamics of subepidermal blister healing are completely unknown. In addition, full-thickness skin wounding, when applied to developmental skin, is unsuitable for distinguishing tissue regeneration and development. In contrast, blistering injury allows us to monitor both skin regeneration (reepithelization of the epidermis) and morphogenesis (HF development) within the same wound bed.

Here, by taking advantage of subepidermal blisters, we explore the effects of injury on developmental tissue. Unexpectedly, blistering injury is found to reduce the expression of tissue morphogenesis genes in the healed epidermis and to direct HFSCs, rather than epidermal SCs, to provide progeny to heal the wound and to suspend HF development.

## Results

### Subepidermal blister formation and its healing process

The suction-blister technique was developed more than a half-century ago to selectively remove the epidermis from the dermis (25, 26), and it has been utilized to harvest epidermal pieces for transplant to repair human skin defects. We reasoned that suction blisters on neonatal mice, in which the epidermis is removed while HFs, the dermis, and subcutaneous fat tissues are maintained in the wounds, enable us to examine the direct relationship between tissue injury and skin development. Therefore, we applied constant negative pressure to the dorsal skin of C57BL/6 wild-type (WT) neonates to produce subepidermal blisters (postnatal day 1 (P1), **Figure 1a**). Histologically, skin separation occurred at the level of the dermoepidermal junction (DEJ) (**Figure 1b**). HFs, as shown by alkaline phosphatase (AP)-positive dermal papillae, remained on the dermal side of the blisters (**Figure 1b**). Dermis and subcutaneous tissues were intact after blistering (**Figure 1b**). α6 integrin (ITGA6), a hemidesmosome protein, was seen at the blister roof, whereas type IV collagen (COL4), a major component of the epidermal basement membrane, and laminin 332 (L332) were present at the base of the blister (**Figure 1c**). In line with the immunofluorescence data, hemidesmosomes localized on the blister roof and lamina densa (basement membrane) were observed at the blister base by electron microscopy (**Figure 1d**), as seen in human suction blisters (25) and their murine counterparts (27).

**Figure 1.**
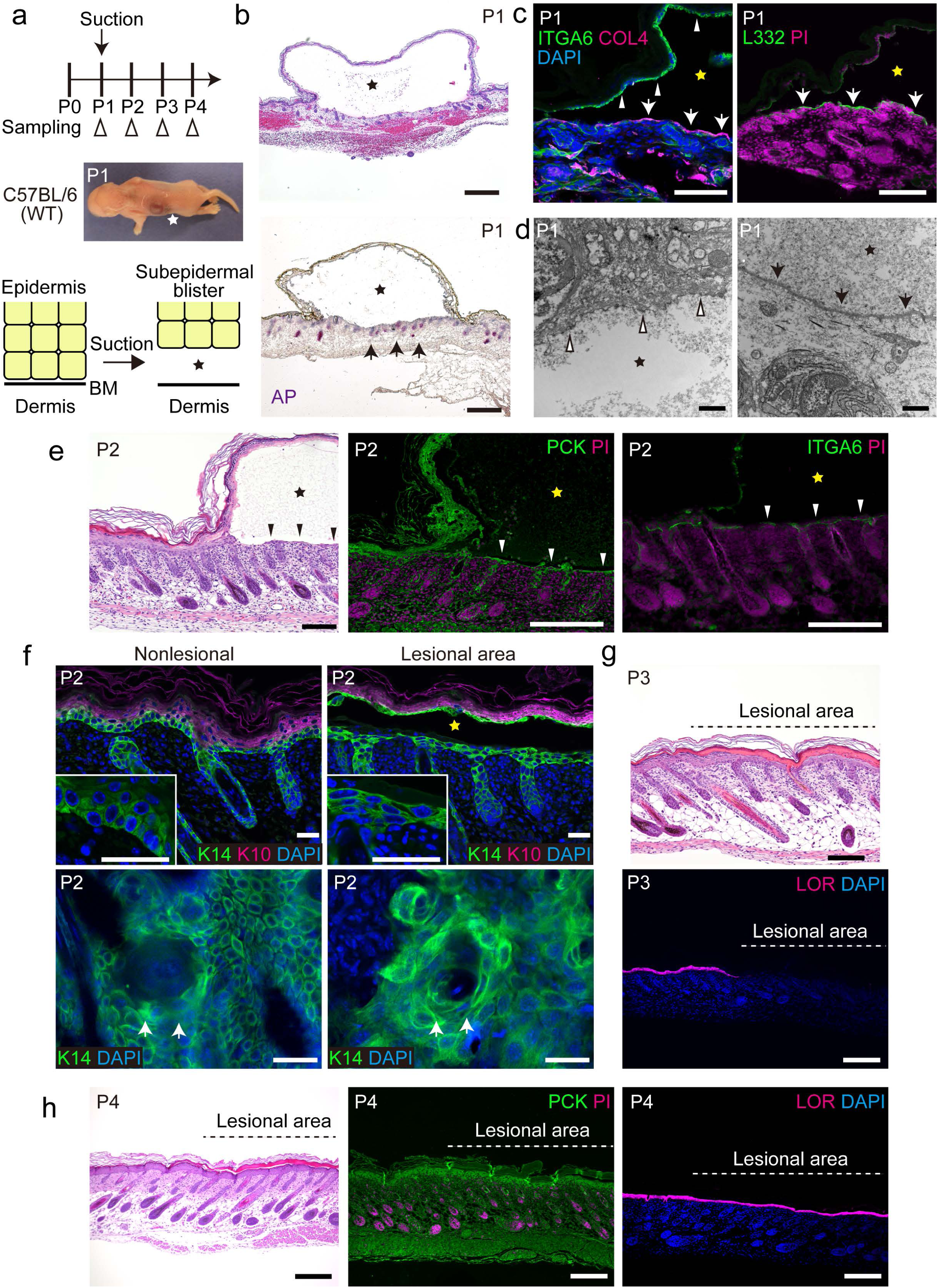
Healing of subepidermal blisters in neonatal mice. (a) Schematic diagram of suction blistering and sample collection. A blister produced on C57BL/6 wild-type (WT) mouse dorsal skin at P1. BM: basement membrane. (b-d) Blistered samples at P1. Hematoxylin and eosin (H&E) and alkaline phosphatase (AP) staining (b). Hair follicles (HFs) remaining in the dermis (indicated by arrows). Scale bar: 500 μm. α6 Integrin (ITGA6, indicated by arrowheads) and type IV collagen (COL4, arrows) labeling (c, left). Laminin 332 (L332, arrows) staining (c, right). Scale bar: 100 μm. Ultrastructural findings of blistered skin (left image: blister roof, right image: blister bottom) (d). Hemidesmosomes (white arrowheads) and lamina densa (arrows) are indicated. Scale bar: 1 μm. (e) H&E, pan-cytokeratin (PCK), and ITGA6 staining at P2. The regenerated epidermis is indicated by arrowheads. Scale bar: 200 μm. (f) Keratin 14 (K14) and keratin 10 (K10) staining of the nonlesional (intact) and lesional (blistered) skin at P2 (upper images and inlets: sections, lower images: whole-mount imaging). HFs are indicated by arrows in the whole-mount images. Scale bar: 30 μm. (g) H&E and loricrin (LOR) staining at P3. Scale bar: 200 μm. (h) H&E, PCK, and LOR staining at P4. Scale bar: 200 μm. Blisters are indicated by stars.

We then characterized the healing processes of the subepidermal blisters. One to two layers of the regenerated epidermis, marked with pan-cytokeratin, was found one day after blister formation (P2, **Figure 1e**). The regenerated epidermis restored ITGA6 expression at the DEJ (**Figure 1e**). The shape of the basal keratinocytes in the intact skin was cuboidal or columnar (P2, keratin 14 (K14)-positive cells in the nonlesional area, **Figure 1f**). In contrast, the regenerated keratinocytes in the blistered skin transformed from cuboidal to a wedge/flattened shape (P2, K14-positive cells in the lesional area, **Figure 1f**) (27). Two days after blistering (P3), the stratified epidermal layers were mostly restored but still lacked loricrin-positive granular layers, a hallmark of proper epidermal differentiation, in the lesional area (**Figure 1g**). The final step in epidermal differentiation was completed by the formation of loricrin-positive granular layers three days after blister formation (P4, **Figure 1h**). These results demonstrate that subepidermal blisters on neonates can serve a model for visualizing wound healing without damaging HFs and other dermal components at the developmental stage.

### Epidermal restoration at the expense of skin development

To elucidate the effects of the blistering injury on the neonatal skin, we performed RNA-seq profiling of the regenerated epidermis one day after blistering (P2, **Figure 2a**). Unexpectedly, the expression of genes involved in HF morphogenesis, such as Wnt signaling, melanogenesis, and Hedgehog signaling, was downregulated (**Figure 2b, 2c**), suggesting that epidermal wounding tunes down tissue development to accelerate blister healing. In agreement with this hypothesis, the number of hair canals, which are present in only developed HFs (HF morphogenesis, stage 6-8) (28), was reduced in the regenerated epidermis (**Figure 2d, 2e**). HF growth under the regenerated epidermis at P4 was delayed at stages 5 and 6, whereas the surrounding intact skin of the blisters or the normal skin of the littermate controls had stage 7 HFs (**Figure 2f**). In Wnt-reporter mice (ins-Topgal+), the LacZ-positive area that is indicative of active Wnt signaling was diminished upon blister formation, accompanied by smaller HFs than those in the surrounding intact skin (P2, **Figure 2g**).

**Figure 2.**
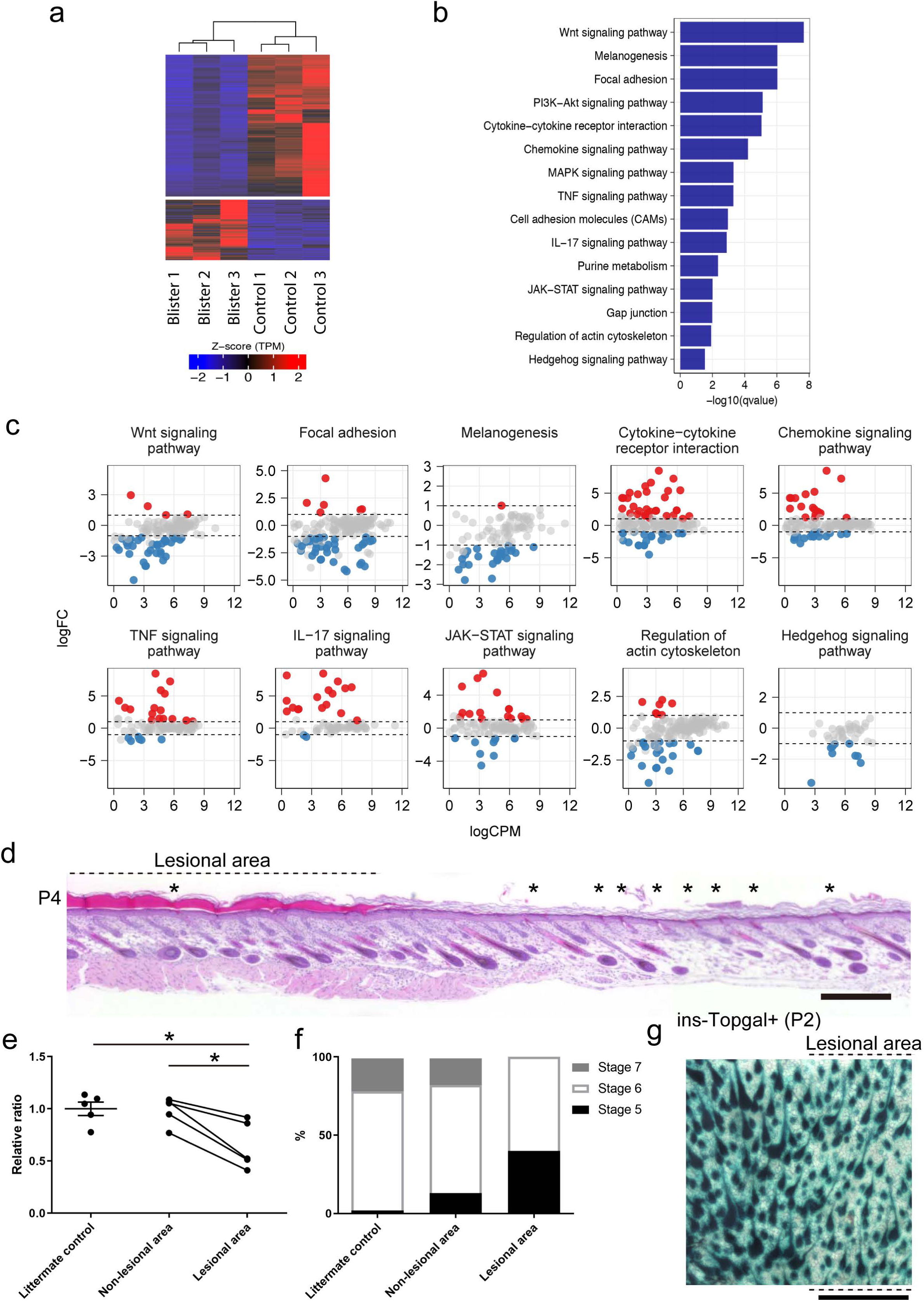
Delayed HF growth during subepidermal blister healing. (a) Heat map (Pearson’s correlation) of differentially expressed genes between the blistered (regenerated) and control WT dorsal skin epidermis at P2 (n=3). (b) GO analysis of differentially expressed genes in the regenerated epidermis. (c) Scatter plots of differentially expressed genes in the regenerated epidermis. (d) Hair canals in the regenerated (lesional) and nonlesional epidermis at P4 (indicated by asterisks). Scale bar: 300 μm. (e) Quantification of hair canals in the lesional, nonlesional, and unaffected littermate control epidermis at P4 (n=5). The data are shown as the mean ± SE (littermate control) or connected with lines showing individual mice. *0.01<p<0.05, one-way ANOVA test, followed by Tukey’s test. (f) HF morphogenesis stages at P4 in lesional, nonlesional, and unaffected littermate control skin (n=5). (g) Whole-mount imaging of the blistered skin of ins-Topgal+ mice at P2. Scale bar: 500 μm.

The expression of genes involved in cytokine-cytokine receptor interactions and chemokine, TNF, IL-17, and JAK-STAT signaling pathways was increased in the regenerated epidermis (**Figure 2b, 2c**), and these pathways are implicated in the recruitment of immune cells. However, the number of neutrophils, lymphocytes, and macrophages was not increased in the lesional dermis (P2, **Supplementary Figure 1a-c**), in which 1-2 layers of the regenerated epidermis covered the wound one day after blistering, suggesting that the immune cells might play a minimal role in the context of blister healing because of the fast regeneration process (<24 hr).

These data indicate that, in the context of skin morphogenesis where the immune system is not yet fully defined, subepidermal blisters heal at the expense of HF growth.

### Progeny of junctional zone SCs represent the main cellular contribution to blister healing

Wounded lesions require epithelial cell proliferation and migration to restore skin integrity. We then investigated the dynamics of epidermal and HF keratinocytes during blister healing. One day after blister formation (P2), BrdU+ cells were abundant in the HFs and the intact epidermis adjacent to the blisters (**Figure 3a**). Cells positive for α5 integrin (ITGA5), a marker of migrating keratinocytes (23), were seen in the HFs within the lesional area and in the epidermal boundary between the blister and the nonlesional area (epidermal tongue) (**Figure 3b**). As HF growth was delayed in the regenerated epidermis (**Figure 2d-g**) and proliferative cells were abundant in HFs of the lesional area (**Figure 3a**), HF keratinocytes were deduced to participate in epidermal regeneration rather than in HF development.

**Figure 3.**
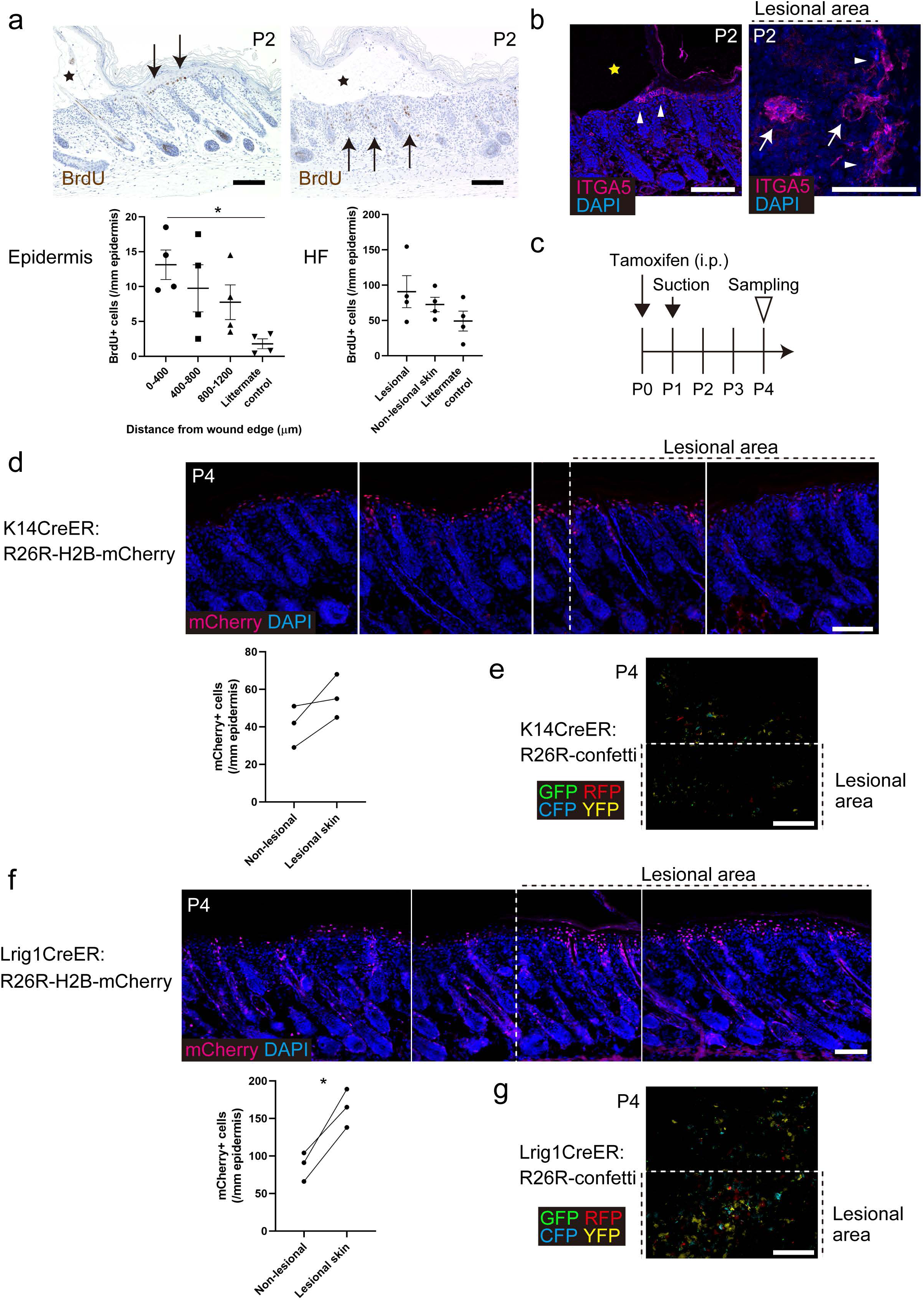
Predominant contribution of HF-derived keratinocytes to subepidermal blister healing. (a) BrdU labeling of blistered samples at P2. Scale bar: 100 μm. BrdU-positive cells are indicated by arrows. Quantification of BrdU-positive cells in the epidermis and HFs (n=4). The data are shown as the mean ± SE. *0.01<p<0.05, one-way ANOVA test, followed by Tukey’s test. (b) α5 integrin (ITGA5) labeling at P2 (left image: section, right image: whole-mount). Scale bar: 100 μm. Blister edges (epidermal tongue) and HFs are indicated by arrowheads and arrows, respectively. Blisters are indicated by stars. (c) Lineage tracing strategy. (d) Sections of K14CreER:H2B-mCherry mice at P4. Quantification of mCherry-positive cells (n=3). Scale bar: 100 μm. The data from individual mice are connected by lines. Student’s t-test. (e) Whole-mount imaging of K14CreER:R26R-confetti samples at P4. Scale bar: 200 μm. (f) Sections of Lrig1CreER:H2B-mCherry mouse skin at P4. Quantification of mCherry-positive cells (n=3). Scale bar: 100 μm. The data from individual mice are connected by lines. *0.01<p<0.05, Student’s t-test. (g) Whole-mount imaging of Lrig1CreER:R26R-confetti mouse samples at P4. Scale bar: 200 μm.

To confirm this hypothesis, we employed a short-term lineage tracing strategy with suction blistering (**Figure 3c**). K14-lineage labeled cells (K14CreER:R26R-H2B-mCherry or K14CreER:R26R-confetti), mainly progeny of SCs in the interfollicular epidermis (IFE), were sparse in the regenerated epidermis (**Figure 3d, 3e**). In contrast, most of the cells in the regenerated epidermis were Lrig1 (leucine-rich repeat and immunoglobulin-like domain protein 1)-lineage labeled cells (Lrig1CreER:R26R-H2B-mCherry or Lrig1CreER:R26R-confetti), which are the progeny of junctional zone SCs (**Figure 3f, 3g**). These data indicate that the HF junctional zone on the dermal side of the blister is the main pool for the keratinocytes that heal subepidermal blisters while halting HF development.

### Depletion of HF junctional stem cells from the wound bed of the blisters promotes the contribution of IFESC progeny to blister healing

The contribution of junctional zone HFSC progeny to blister healing led us to investigate how the epidermis regenerates in the absence of HFs at the blister base (dermis). Type XVII collagen (COL17) is expressed not only in the IFE but also in the bulge region of the HFs (**Figure 4a**) (11, 29-32). COL17 is encoded by the *COL17A1* gene, and its deficiency leads to junctional EB (33). The splitting of neonatal *Col17a1*^-/-^ (34) dorsal skin upon suction blistering was observed between ITGA6 and COL4/L332 (**Figure 4b, 4c**), as was the case in wild-type neonates (**Figure 1c, 1d**). Intriguingly, suction blistering (**Figure 1a**) of *Col17a1*^-/-^ dorsal skin detached HFs from the dermis (P1, **Figure 4d**). In agreement with this finding, dermal papilla cells (AP+) were observed on the roof side of the blisters of *Col17a1*^-/-^ mice, whereas the blister roofs of control mice did not have these cells (P1, **Figure 4e**). Epidermal regeneration was not apparent in *Col17a1*^-/-^ mice one day after suction blistering (P2), whereas the control mice showed regeneration of the epithelial layers (**Figure 4f**). *Col17a1*^-/-^ mice had delayed expression of loricrin in the regenerated epidermis three days after blister formation (P4, **Figure 4f**). BrdU+ cells were abundant in the *Col17a1*^-/-^ mouse epidermis surrounding blisters as was the case for controls (**Figures 3a, 4g**). Lineage tracing experiments (**Figure 3c**) revealed that IFESC progeny covered most of the regenerated area (K14CreER:R26R-H2B-mCherry:*Col17a1*^-/-^) at P4 (**Figure 4h, 4i**). The transgenic rescue of *Col17a1*^-/-^ by overexpressing human COL17 (hCOL17+;*Col17a1*^-/-^) (34) normalized blister healing (P4, **Supplementary Figure 2**). These data demonstrate that upon HF depletion, the IFE can compensate the lack of junctional stem cells and repair defects in the epidermis.

**Figure 4.**
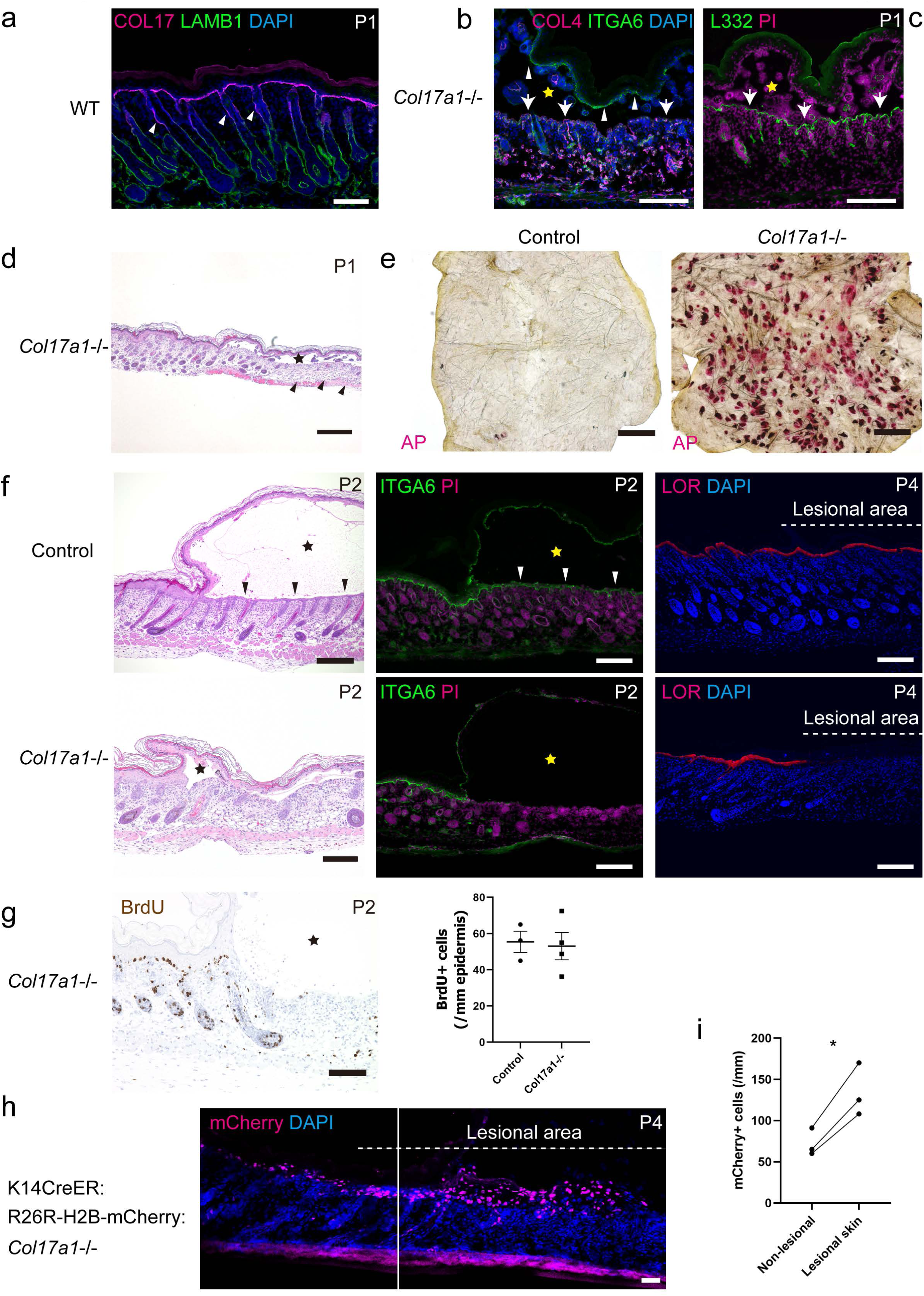
Effects of HF depletion on subepidermal blister healing. (a) Type XVII collagen (COL17, arrowheads indicate the hair bulge) and laminin β1 (LAMB1) labeling in WT dorsal skin sections (P1). Scale bar: 100 μm. (b, c) Blistered samples of *Col17a1*^-/-^ mouse dorsal skin at P1. ITGA6 (indicated by arrowheads) and COL4 (arrows) labeling (b). L332 staining (c, arrows). Scale bar: 100 μm. (d) H&E staining of blistered skin from *Col17a1*^-/-^ mice at P1. HFs detached from the dermis in *Col17a1*^-/-^ skin are indicated by arrowheads. Scale bar: 500 μm. (e) Whole-mount AP staining of the blister roof epidermis from *Col17a1*^-/-^ mice and littermate controls at P1. Scale bar: 500 μm. (f) H&E (P2), ITGA6 (P2) and LOR staining (P4) of *Col17a1*^-/-^ mice and littermate controls. The regenerated epidermis is indicated by arrowheads. Scale bar: 200 μm. (g) BrdU labeling of *Col17a1*^-/-^ skin at P2. Scale bar: 100 μm. Quantification of BrdU-positive cells in the epidermis surrounding blisters (n=4). The data are shown as the mean ± SE. Student’s t-test. (h, i) Lineage tracing of K14CreER:R26R-mCherry:*Col17a1*^-/-^ at P4 (h). Scale bar: 100 μm. Quantification of mCherry-positive cells in the regenerated epidermis (n=3) (i). The data from individual mice are connected by lines. *0.01<p<0.05, Student’s t-test. Blisters are indicated by stars.

### Impaired flattening of regenerated keratinocytes delays blister healing

We further sought to identify other modulators of subepidermal blister healing. We first focused on collagen VII (COL7), encoded by *Col7a1*. COL7 forms anchoring fibrils and is located at the DEJ (**Figure 5a**) but just below the basement membrane (35, 36), and its deficiency leads to dystrophic EB (37, 38). As conventional wound healing is delayed in COL7-hypomorphic mice (39), we applied the suction-blister method to *Col7a1*^-/-^ mice (40) (**Figure 5b-h, Supplementary Figure 3**). In contrast to that in WT and *Col17a1*^-/-^ suction blisters (**Figure 1c, 1d, 4b, 4c**), skin splitting occurred at the level below the basement membrane in *Col7a1*^-/-^ mouse dorsal skin, as shown by the presence of L332 and COL4 on the blister roof epidermis (P1, **Figure 5b, 5c**). The epidermal defects were not repaired in *Col7a1*^-/-^ mice, whereas the epidermis of the control blistered skin regenerated one day after blistering (P2, **Figure 5f, Supplementary Figure 3**), which is consistent with the delayed healing of full-thickness skin wounds in COL7-hypomorphic mice (39). This finding is contrasted by the fact that COL7-depleted keratinocytes migrate faster than WT keratinocytes in vitro (41, 42). We examined the HFs of *Col7a1*^-/-^ mice to explain the slowed epidermal regeneration because HFs are the main contributor to blister healing (**Figure 2a-g**). However, HFs were present in the *Col7a1*^-/-^ mouse wound bed (blister base) (P1, **Figure 5d, 5e**) as opposed to that of *Col17a1*^-/-^ mouse (P1, **Figure 4d, 4e**). Moreover, the number of BrdU+ cells in HFs was comparable between *Col7a1*^-/-^ and control mice (P2, **Figure 5g, Supplementary Figure 3**), suggesting that the proliferation of HF keratinocytes does not account for the delayed blister healing of *Col7a1*^-/-^ mice.

**Figure 5.**
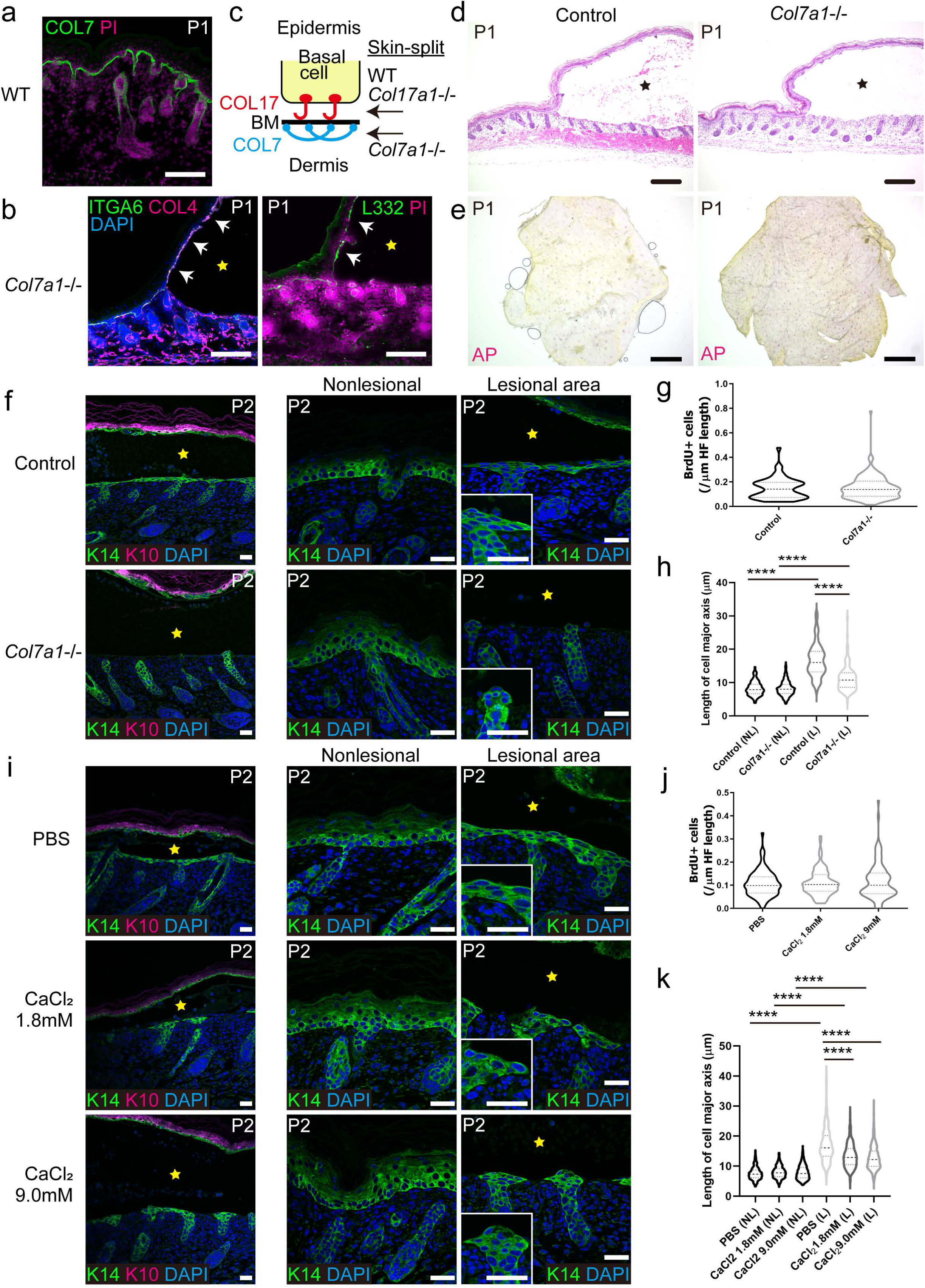
Involvement of keratinocyte shape transformation in subepidermal blister healing. (a) Type VII collagen (COL7) labeling in WT dorsal skin sections (P1). Scale bar: 200 μm. (b) ITGA6/COL4 (left, indicated by arrows) and L332 labeling (right, indicated by arrows) in the blistered skin of *Col7a1*^-/-^ mice at P1. Scale bar: 100 μm. (c) Schematic diagram of control, *Col17a1*^-/-^, and *Col7a1*^-/-^ mouse skin splits. BM: basement membrane. (d) H&E staining of blistered skin from *Col7a1*^-/-^ mice and their littermate controls at P1. Scale bar: 200 μm. (e) Whole-mount AP staining of the blister roof epidermis from *Col7a1*^-/-^ mice and their littermate controls at P1. Scale bar: 500 μm. (f) K10/K14 (low magnification) and K14 (high magnification) labeling of *Col7a1*^-/-^ mouse and littermate control blistered skin at P2. Scale bar: 30 μm. (g) Quantification of BrdU-positive cells per μm HF length (n=55 HFs from three control and 143 HFs from four *Col7a1*^-/-^ mice). The data are shown as violin plots. Student’s t-test. (h) Length of the major axis of keratinocytes in the regenerated epidermis (n=244 (control, L), and 132 (*Col7a1*^-/-^, L) cells from four mice, respectively) and in the surrounding intact epidermis (basal cells; n=200 (control, NL), 299 (*Col7a1*^-/-^, NL), from four mice, respectively). NL: nonlesional area. L: lesional area. The data are shown as violin plots. ****p<0.0001, one-way ANOVA test, followed by Tukey’s test. (i) K10/K14 and K14 (high magnification) labeling of WT blistered skin treated with CaCl_2_ or PBS at P2. Scale bar: 30 μm. (j) Quantification of BrdU-positive cells per μm HF length (n=83 (PBS), 95 (1.8 mM CaCl_2_), and 97 (9.0 mM CaCl_2_) HFs from four mice). (k) Length of the major axis of keratinocytes in the regenerated epidermis (n=433 (PBS, L), 451 (1.8 mM CaCl_2,_ L), and 425 (9.0 mM CaCl_2_, L) cells from four mice) and in the surrounding intact epidermis (basal cells; n=311 (PBS, NL), 279 (1.8 mM CaCl_2,_ NL), 302 (9.0 mM CaCl_2_, NL) cells from four mice). NL: nonlesional area. L: lesional area. The data are shown as violin plots. ****p<0.0001, one-way ANOVA test, followed by Tukey’s test. Blisters are indicated by stars.

Second, we treated the blistered skin in wild-type mice with extracellular calcium (**Figure 5i-k, Supplementary Figure 3**). Extracellular calcium is a potent inhibitor of proliferation and migration in cultured keratinocytes as well as an inducer of differentiation (43, 44). Consistent with previous in vitro assays, the intrablister administration of CaCl_2_ (1.8 mM or 9.0 mM) just after suction blistering delayed epidermal regeneration in vivo (P2, **Figure 5i**). Premature differentiation, which might hinder wound healing, was not apparent in the CaCl_2_-treated blisters, as K10 labeling was seen only at the blister roof but not in the keratinocytes on the wound bed (P2, **Figure 5i**). Similar to that in *Col7a1*^-/-^ mouse blisters, the number of BrdU+ cells in HFs was not reduced in CaCl_2_-treated blisters (P2, **Figure 5j, Supplementary Figure 3**).

These two examples strongly suggest that there are factors other than HF keratinocyte proliferation that modulate blister healing. During blister healing, keratinocytes reshape into a wedge-shaped morphology (**Figure 1f**), which is mirrored by the RNA-seq data showing that the expression of genes involved in the regulation of the actin cytoskeleton is decreased in the regenerated epidermis (**Figure 2b, 2c**). Wedge-shaped/flattened keratinocytes are believed to be superior to cuboidal/columnar keratinocytes for covering epidermal defects. In the intact (nonblistered) skin, the morphology of *Col7a1*^-/-^ or Ca-treated basal keratinocytes was similar to that of control cells (P2, the nonlesional area in **Figure 5f, 5h, 5i, 5k**). The regenerated keratinocytes became wedge-shaped/flattened in the control group, as shown in **Figure 1f**. However, the keratinocytes in the regenerated epidermis of *Col7a1*^-/-^ or Ca-treated mice were not as flat as those of control mice but were still rather cuboidal (P2, the lesional area in **Figure 5f, 5h, 5i, 5k**), which, at least partly, explains the delayed blister healing in these mice. These data imply that morphological changes in keratinocytes contribute to subepidermal blister healing.

### Mathematical modeling reproduces blister healing

These in vivo experiments led us to speculate that HF/IFE cell proliferation during paused HF development and the morphological changes of the regenerated keratinocytes might simply account for the dynamics of subepidermal blister healing. To answer this question, we employed mathematical modeling. The particle-based model of self-replicating cells (45, 46) allowed us to visualize the dynamics of the epidermal basal layer and to establish the epidermal defects on the basement membrane. We utilized the data on the number of BrdU+ cells among HF vs. IFE keratinocytes (approximately 7:1 per unit of epidermal length; **Figure 3a**) and on the shape of the regenerated vs. normal keratinocytes (2:1 the length of the major cell axis; **Figure 5h, 5k**). SC progeny within the epidermal defects (colored in red), simulating HF-derived cells, had a more substantial contribution to wound healing than IFE SCs (colored in yellow; **Figure 6a, 6b, Supplementary Video 1**), as seen in the lineage tracing experiments (**Figure 3c-g**). The absence of SCs within the epidermal defects, simulating HF depletion in the wound bed, showed delayed healing (**Figure 6c, 6d, Supplementary Video 2**), in agreement with *Col17a1*^-/-^ epidermal regeneration results (**Figure 4e-g**). A less flattened morphology of the regenerated keratinocytes, simulating *Col7a1*^-/-^ and Ca-treated blister healing (regenerated vs. normal keratinocytes, 1.3-1.6:1 in length; **Figure 5h, 5k**), slowed epidermal regeneration (**Figure 6e, 6f, Supplementary Video 3**). These in silico data demonstrate that the contribution of HFSC progeny and the morphological change in the regenerated keratinocytes are sufficient to recapitulate the in vivo subepidermal blister healing.

**Figure 6.**
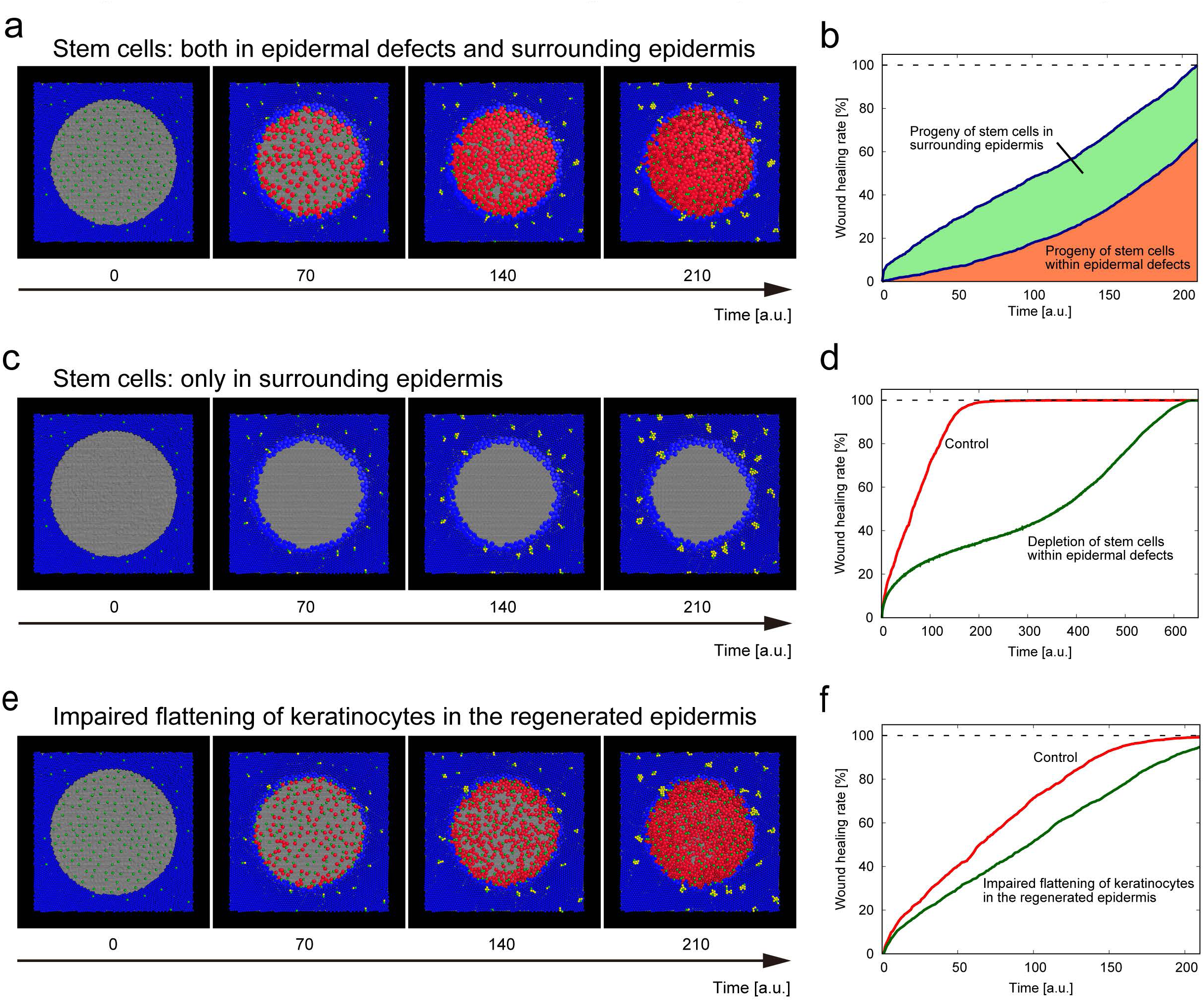
Mathematical modeling of subepidermal blister healing. (a) A particle-based model of subepidermal blister healing at the basal layer. Epidermal basal cells (colored in blue), which do not divide, are placed on the basement membrane (gray). Stem cells (SCs, green) give rise to progeny (simulating HF-derived cells; red) within epidermal defects or in the surrounding epidermis (IFE-derived cells; yellow). t: arbitrary time. See **Supplementary Video 1**. (b) Contribution of each progeny cell within the epidermal defect or of surrounding epidermis to subepidermal blister healing, measured as the ratio of the area occupied by each progeny to the area of the initial epidermal defect. (c) A model of subepidermal blister healing without SCs within epidermal defects. See **Supplementary Video 2**. (d) Time course of subepidermal blister healing in control (a) and SC-depleted epidermal defects (c). (e) Effects of the impaired flattening of keratinocytes upon epidermal regeneration. The diameter of basal keratinocytes (long axis of the spheroid) in the regenerated vs. surrounding epidermis was calculated as 1.5:1 (in contrast to 2:1 in Figure 6a). See **Supplementary Video 3**. (f) Time course of wound healing for control (a) and less flattened keratinocytes in the regenerated epidermis (e).

We finally asked whether this HF contribution to wound healing could be applied to a human setting, in which HFs are larger but much more sparsely distributed than their murine counterparts (**Supplementary Table 1**). Mathematical modeling revealed that the progeny of HF keratinocytes are still the main contributors to subepidermal blister healing in humans (**Supplementary Figure 4a, b, Supplementary Video 4**). To further confirm the in silico modeling results, we examined human subepidermal blister samples and found that epidermal regeneration by HFs was observed in the samples with the re-epithelized area (**Supplementary Figure 4c**). These findings strongly suggest that HFs can replenish keratinocytes for subepidermal blister healing in both mice and humans.

## Discussion

Although recent studies have reported that injury causes damaged tissue to shift to an embryonic-like state, there is a poor understanding of how regeneration affects development at the damaged tissues. Here, we applied the blistering injury to neonatal mouse dorsal skin and showed the skewed contribution of HFSC progeny to wound healing rather than HF development.

Previous studies on skin wounding combined with the fate mapping of murine skin delineated the involvement of epithelial, mesenchymal, and immune cells in wound healing, depending on different settings (12, 13). However, as full-thickness skin wounds, even when they are applied to neonatal skin, remove all skin components, it is challenging to see the effects of development on injury or vice versa. Our blistering injury has an advantage over conventional skin wounding studies in that only the epidermis is removed by constant negative pressure, with the other skin components and basement membrane being retained in the wounds, which allowed us to examine “pure” epidermal wound healing processes during skin development. Our study contrasts with wound-induced embryonic gene expression (1) and follicular neogenesis upon full-thickness skin wounding in adult (47, 48) and neonatal mice (49). It is noteworthy that subepidermal blister healing suppressed Wnt signaling in our study (**Figure 2a-2g**), whereas Wnt activation has been implicated in full-thickness skin wounds (48), implying context-dependent Wnt activities in tissue regeneration.

Previous studies have suggested the possible involvement of HFs in epidermal regeneration in human suction blisters (50) and extracellular matrix alterations in the skin-split area (51, 52). We show that the progeny of HF junctional zone SCs mainly repair subepidermal blisters (**Figures 3f, 3g, 6a, 6b**). However, the IFE can also serve as a reservoir of keratinocytes to repair epidermal defects when HFs are depleted (**Figure 4h**). How the contribution of two sources of keratinocytes to blister healing is regulated is unknown, but the significant contribution of HF junctional zone progeny is reasonable because HFs are densely located in the wound bed (blister base). In contrast, the progeny of IFESCs could recover only from the blister edge, as demonstrated by mathematical modeling (**Figures 3d-3g, 6a, 6b**). The role of HFSCs is also highlighted by delayed HF growth in the regenerated epidermis, as corroborated by the downregulated Wnt signaling (**Figure 2a-2g**). These findings indicate that there is a coordinated balance between tissue development and wound healing, which has not been well recognized. In addition, the expression of IL-17 signaling pathway genes was increased, although the recruitment of immune cells was not evident one day after blistering (**Supplementary Figure 1**). Recently, IL-17 signaling has been shown to drive Lrig1-lineage cell recruitment in wound healing and tumorigenesis (53). Therefore, IL-17 signaling might also help Lrig1-lineage cells translocate from HFs to repair epidermal defects in our study (**Figure 3f, 3g**).

The dynamics of cytoskeletal changes directly affect cellular morphology and migration potential (54). Previous studies have shown that cells undergo a morphological transformation into a wedge/flattened shape at the leading edge of migrating cells (55) and in regenerated keratinocytes during wound healing (27, 56). Our study has shed further light on the significant impact of the keratinocyte morphological changes on in vivo wound healing through blistering experiments in *Col7a1*^-/-^ and Ca-treated mice and mathematical modeling (**Figure 5f, 5h, 5i, 5k, 6a, 6e, 6f**).

Our in vivo blistering experiments can mimic and replace the in vitro cultured cell wound healing assay (e.g., scratch wounding), which has been used in the field of cell biology for decades because subepidermal blisters are epidermal wounds and the involvement of mesenchymal and immune cells are not pivotal for blister healing. In vivo intrablister administration of drugs, as exemplified by extracellular calcium administration (**Figure 5i**), could be an alternative to in vitro chemical treatment of scratch-wounded cultured cells to develop new therapeutic options for wound healing, especially of subepidermal blisters.

Our in vivo suction-blister model recapitulates the human pathological epidermal detachment seen in EB, pemphigoid diseases, burns, and severe drug reactions such as Stevens-Johnson syndrome/toxic epidermal necrolysis. Loss-of-function mutations in *COL17A1* (33) and *COL7A1* (37, 38) lead to junctional and recessive dystrophic EB in humans, respectively. Therefore, *Col17a1*^-/-^ and *Col7a1*^-/-^ mouse blistering also serves as an EB wound model. The prominent hair loss in human *COL17A1*-mutated junctional EB might be reflected by the depletion of HFs from the wound bed in *Col17a1*^-/-^ blisters (**Figure 4d, 4e**), whereas the hair loss in recessive dystrophic EB is not as severe as that in the junctional subtype (57), consistent with the maintenance of HFs in the dermis of *Col7a1*^-/-^ mouse dorsal skin upon suction blistering (**Figure 5d, 5e**). Furthermore, the processes of subepidermal blister healing highlight HFs as a target for treating the wounds of EB and other blistering diseases.

In closing, our study has revealed the imbalance between development and wound regeneration in the skin blisters. In contrast to wound healing in adult skin, where embryonic programs are reactivated, blister repair during development temporarily inhibits the Wnt pathway, a crucial skin morphogenic driver, leading to the biased activities of HF junctional stem cells from HF morphogenesis towards wound healing. Our findings of the healing processes pave the way for tailored therapeutic interventions for epidermolysis bullosa, pemphigoid diseases and other blistering diseases.

## Material and Methods

### Animals

C57BL/6 strain mice were purchased from Clea (Tokyo, Japan). Ins-Topgal+ mice were obtained from RIKEN BRC (Tsukuba, Japan) (58). K14CreER, Lrig1CreER, and R26R-confetti mice were purchased from the Jackson Laboratory (Bar Harbor, Maine, USA). R26R-H2B-mCherry mice were provided by RIKEN (Kobe, Japan). *Col17a1*^-/-^ and hCOL17+;*Col17a1*^-/-^ mice were generated as previously described (34). *Col7a1*^-/-^ mice were provided by Prof. Jouni Uitto (40). The institutional review board of the Hokkaido University Graduate School of Medicine approved all animal studies described below.

### Suction blisters

Suction blisters were produced on the neonatal murine dorsal skin (P1) using a syringe and connector tubes. The negative pressure applied to the skin (generally for minutes) was 523.4±1.3 mmHg (evaluated by an Ex Pocket Pressure Indicator PM-281 (AS ONE, Osaka, Japan)). The diameter of the syringe attached to the skin was 4 mm. The size of the typical blister was 3 mm in diameter.

### Histology

Mouse dorsal skin specimens were fixed in formalin and embedded in paraffin after dehydration or were frozen on dry ice in an optimal cutting temperature (OCT) compound. Frozen sections were fixed with 4% paraformaldehyde (PFA) or cold acetone or were stained without fixation. Antigen retrieval with pH 6.0 (citrate) or pH 9.0 (EDTA) buffer was performed on deparaffinized sections. Sections were incubated with primary antibodies overnight at 4°C. After being washed in phosphate-buffered saline (PBS), the sections were incubated with secondary antibodies conjugated to FITC, Alexa 488, Alexa 647 or Alexa 680 for 1 hr at room temperature (RT). The nuclei were stained with propidium iodide (PI) or 4’,6-diamidino-2-phenylindole (DAPI). The stained immunofluorescent samples were observed using a confocal laser scanning microscope (FV-1000 (Olympus, Tokyo, Japan) or LSM-710 (Zeiss, Oberkochen, Germany)).

For immunohistochemistry, horseradish peroxidase (HRP)-tagged secondary antibodies were used. Sections were blocked with hydrogen peroxide, labeled with antibodies, and counterstained with hematoxylin. For morphological analysis, deparaffinized sections were stained with hematoxylin and eosin (H&E) by conventional methods. Alkaline phosphatase staining was performed using a StemAb Alkaline Phosphatase Staining Kit II (Stemgent, San Diego, California, USA). Images of immunohistochemistry, and H&E- and alkaline phosphatase-stained sections were captured with a BZ-9000 microscope (Keyence, Tokyo, Japan).

For whole-mount staining, mouse dorsal skin samples were fixed with 4% PFA and immunolabeled or stained with the Alkaline Phosphatase Staining Kit II. For X-gal staining of ins-Topgal+ mouse skin, a beta-galactosidase staining kit (Takara-bio, Shiga, Japan) was used according to the provider’s protocol. Briefly, dorsal skin samples were fixed with 4% PFA for 1 hr at 4°C and soaked in staining solution overnight at RT. Tissues were mounted in a Mowiol solution. Images were observed with LSM-710, FV-1000 or BZ-9000 microscopes.

HF morphological stages were evaluated as previously described (28). The length of the major axis of keratinocytes in the intact and regenerated epidermis was measured using ImageJ (NIH, Bethesda, Maryland, USA) on K14-stained sections. The quantification of the cells expressing a particular marker was performed as previously described (59).

### Antibodies

The following antibodies were used: anti-BrdU (Abcam, Cambridge, UK; BU1/75, Dako; M0744), anti-loricrin (Covance, Princeton, New Jersey, USA), FITC-conjugated anti-CD3e (BioLegend, San Diego, California, USA; 145-2C11), Alexa Fluor 488-conjugated anti-F4/80 (Affymetrix, Santa Clara, California, USA; BM8), FITC (fluorescein isothiocyanate)-conjugated anti-Ly-6G (Beckman Coulter, Brea, California, USA; RB6-8C5), anti-COL4 (Novus Biologicals, Centennial, Colorado; NB120-6586), anti-COL7 (homemade (60)), anti-COL17 (Abcam; ab186415), anti-ITGA5 (Abcam; EPR7854), anti-ITGA6 (BD Biosciences Pharmingen, San Diego, California, USA; GoH3), anti-L332 (Abcam; ab14509), anti-laminin β1 (Abcam; ab44941), anti-pan-cytokeratin (PROGEN, Wieblingen, Heidelberg, Germany; PRGN-10550), anti-cytokeratin 10 (Biolegend; Poly19054), anti-cytokeratin 14 (ThermoFisher, Waltham, Massachusetts, USA; LL002).

### BrdU labeling

For proliferation analysis, 10 μg of BrdU (BD Biosciences Pharmingen) per head was intraperitoneally administered 4 hr before sacrifice.

### Transmission electron microscopy

The samples were taken from C57BL/6 mouse dorsal skin (P1) just after suction blistering was performed. The samples were fixed in 5% glutaraldehyde solution, postfixed in 1% OsO_4_, dehydrated, and embedded in Epon 812. The embedded samples were sectioned at 1 µm thickness for light microscopy and thin-sectioned for electron microscopy (70 nm thick). The thin sections were stained with uranyl acetate and lead citrate and examined by transmission electron microscopy (H-7100; Hitachi, Tokyo, Japan).

### Lineage tracing

K14CreER:R26R-H2B-mCherry, Lrig1CreER:R26R-H2B-mCherry, K14CreER:R26R-confetti, and Lrig1CreER:R26R-confetti mice were intraperitoneally treated with 0.5 mg of tamoxifen (T5648; Sigma-Aldrich, St. Louis, Missouri, USA) at P0. The dorsal skin samples were harvested four days later (P4).

### RNA sequencing and analysis

Suction-blistered and control samples were collected at P2 and treated with 0.25% trypsin EDTA overnight at 4°C. The epidermis was minced with a scalpel and suspended in 10% FCS DMEM. The cell suspension was filtered through a 70 µm filter, and cell pellets were collected. Library preparation was performed using an Illumina TruSeq RNA prep kit by following the manufacturer’s instructions. Briefly, following TRIzol extraction and chemical fragmentation, mRNA was purified with oligo-dT-attached magnetic beads and reverse transcribed into cDNA. Following a second strand synthesis step with DNA polymerase I and RNAse H, the resulting cDNA was subjected to end repair, A-tailing, and Illumina compatible adaptor ligation. Following purification and PCR-mediated enrichment, libraries were purified with AMPure XP beads and sequenced on a NextSeq 500 Illumina sequencer.

After quality controls were performed, the raw reads were aligned to the NCBIm37 mouse reference genome (mm9) using HiSat2 (61) (version 2.0.0) using options -N 1 -L 20 -i S, 1, 0.5 -D 25 -R 5 --pen-noncansplice 20 --mp 1, 0 --sp 3, 0 and providing a list of known splice sites. Expression levels were quantified using featureCounts (62) with RefSeq gene annotation and normalized as TPM using custom scripts. Differential expression analysis was performed using the edgeR (63) software package. After lowly expressed genes (1 count per million in less than two samples) were filtered out, the treatment and control groups were compared using the exact test method (63). Genes with an absolute log2-fold change greater than 1 and false discovery rate (FDR) less than or equal to 0.05 were considered differentially expressed. Hierarchical clustering of gene expression profiles was performed on differentially expressed genes using only Euclidean distances and the complete linkage method. TPM values were normalized as Z-scores across samples, and the distances were computed. GO term and KEGG enrichment analysis on the differentially expressed genes were performed with Enrich (64).

### Intrablister administration

Ten microliters of 1.8 or 9.0 mM CaCl_2_ in PBS was administered by syringe into the blisters just after the suction blistering procedure was performed.

### Statistics

Statistical analyses were performed using GraphPad Prism (GraphPad Software, La Jolla, California, USA). P-values were determined using Student’s t-test or one-way ANOVA followed by Tukey’s test. P-values are indicated as *0.01<p<0.05, **0.001<p<0.01, ***0.0001<p<0.001, and ****p<0.0001. The values were shown as the means ± standard errors (SE), violin plots or connected with lines showing individual mice.

### Mathematical modeling

A mathematical model proposed for epidermal cell dynamics (45) was adapted to simulate epidermal wound healing. In this model, epidermal basal cells were represented as spherical particles, with the cell diameter set to 10 µm. Cells designated as epidermal SCs and their progeny could undergo division on the basement membrane. Cell division was described as a process of two initially completely overlapping particles gradually separating into two distinct particles. When a newly created cell was not fully surrounded by other cells, it was judged as being in a regeneration process and immediately underwent a transition to an oblate spheroid shape with the long axis increased by a factor of 2 (normal) or 1.5 (simulating *Col7a1*^-/-^ and Ca-treated blisters) while its volume was kept constant. The same division rate was assigned to all proliferative cells, with an average division period of 57.6 [arb. unit]. Forces exerted on a cell came from adhesion and excluded-volume interactions with other cells and with the basement membrane. SCs were tightly bound and unable to detach from the basement membrane, while the progeny were weakly bound so that they could detach from the membrane via the ambient pressure: the detached cells were removed from the system. The basement membrane was assumed to be a rigid flat surface, whose shape remained unchanged over time. These interactions were calculated to obtain the time evolution of the whole system by solving equations given in a previous report (45). The simulation region was set to 600 µm x 600 µm horizontally with periodic boundary conditions. To prepare the initial conditions for the simulation of subepidermal blister healing, we first ran a simulation with SCs placed on the basement membrane until their progeny covered the whole surface. Then, we set the progeny to be nonproliferative and created epidermal defects by removing the cells that were inside a disk domain with a diameter of 480 µm.

The number of HFs in humans was estimated from the data in Supplementary Table 1 under the assumption that HFs were uniformly distributed and that the number of proliferative cells per HF was proportional to the volume of the HF. Using the mean values of the data, we estimated that the ratio of human to mouse proliferative cells in epidermal defects is 0.174:1. The initial number of cells was prepared accordingly in the simulations of human epidermal defects.

### Human samples

From the H&E-stained skin of patients with congenital or autoimmune subepidermal blistering diseases (73 EB or 188 bullous pemphigoid (BP) samples, respectively), the samples that met the following histological criteria were selected: (1) subepidermal blisters or a skin split at the dermoepidermal junction; (2) re-epithelization in the area; and (3) presence of HFs on the blister base. Three BP samples fulfilled all the criteria (blisters 1, 2, and 3) and were observed with a Keyence BZ-9000 microscope. The institutional review board of the Hokkaido University Graduate School of Medicine approved all human studies described above (ID: 13-043 and 15-052). The study was conducted according to the Declaration of Helsinki Principles. Participants or their legal guardians provided written informed consent.

## Author contributions

Y. F. designed and performed the experiments, analyzed the data, interpreted the results, and wrote the manuscript. M. W., S. T., H. N., H. K., Y. W., Y. M, A. L., V. P., H. U., H. I., W. N., and S. O. performed the experiments and analyzed the data. K. O., Y. K., and M. N. performed the mathematical modeling and wrote the manuscript. G. D. designed and performed the experiments, analyzed the data and interpreted the results. S. H. interpreted the results and supervised the study. K. N. conceived and designed the experiments, analyzed the data, interpreted the results, wrote the manuscript and supervised the study.

## Acknowledgments

We thank Ms. Meari Yoshida for her technical assistance. We also thank Professor Yumiko Saga, Professor Kim B Yancey, and Professor Jouni Uitto for providing the ins-Topgal+, K14-hCOL17, and *Col7a1*^-/-^ mice, respectively. This work was funded by AMED (ID: 18059057), JSPS (KAKEN 17K16317), the Uehara Foundation, the Lydia O’Leary Memorial Pias Dermatological Foundation to KN, JST CREST (JPMJCR15D2) to MN, JSPS (KAKEN 16K10120) to HN, AIRC IG 20240 to SO, and AIRC MFAG 2018 (ID: 21640) to GD.

