## Supplementary Information for "Hair follicle stem cell progeny heal blisters while pausing skin development"

Supplementary Figure 1. Quantification of immune cells in subepidermal blister healing

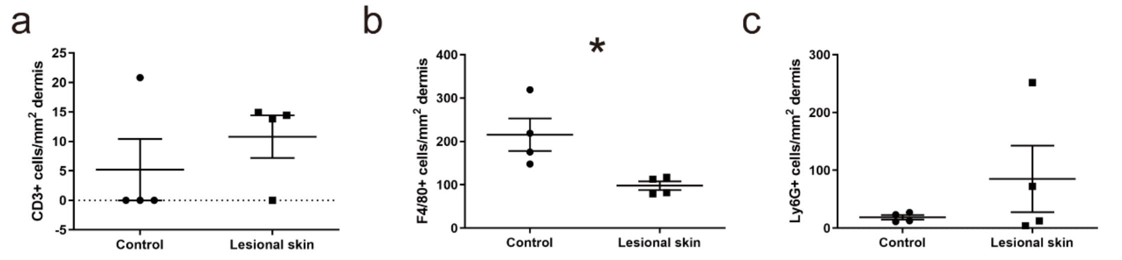


**Supplementary Figure 1. Quantification of immune cells in subepidermal blister healing.**

Quantification of immune cells (CD3 (a), Ly6G (b), and F4-80 (c)) in the dermis of blistered WT and unaffected littermate control skin at P2 (n=4). The data from individual mice are connected by lines. *0.01<p<0.05, Student's t-test.

Supplementary Figure 2.
Transgenic rescue of *Col17a1*-/- blister healing

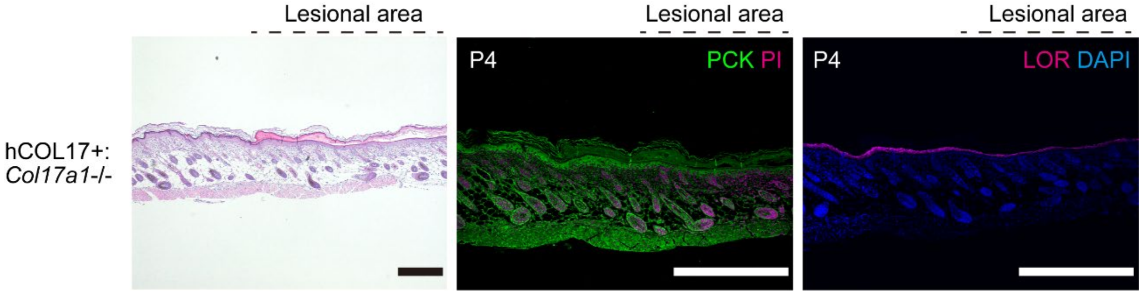


**Supplementary Figure 2. Transgenic rescue of *Col17a1*-/- mouse blister healing.**

Blistered samples of *Col17a1*-/- mice rescued with human COL17 overexpression (hCOL17+;*Col17a1*-/-). H&E, PCK and LOR staining (P4). Scale bar: 500 μm.

Supplementary Figure 3. H&E and BrdU labeling of *Col7a1*-/- and Ca-treated blisters

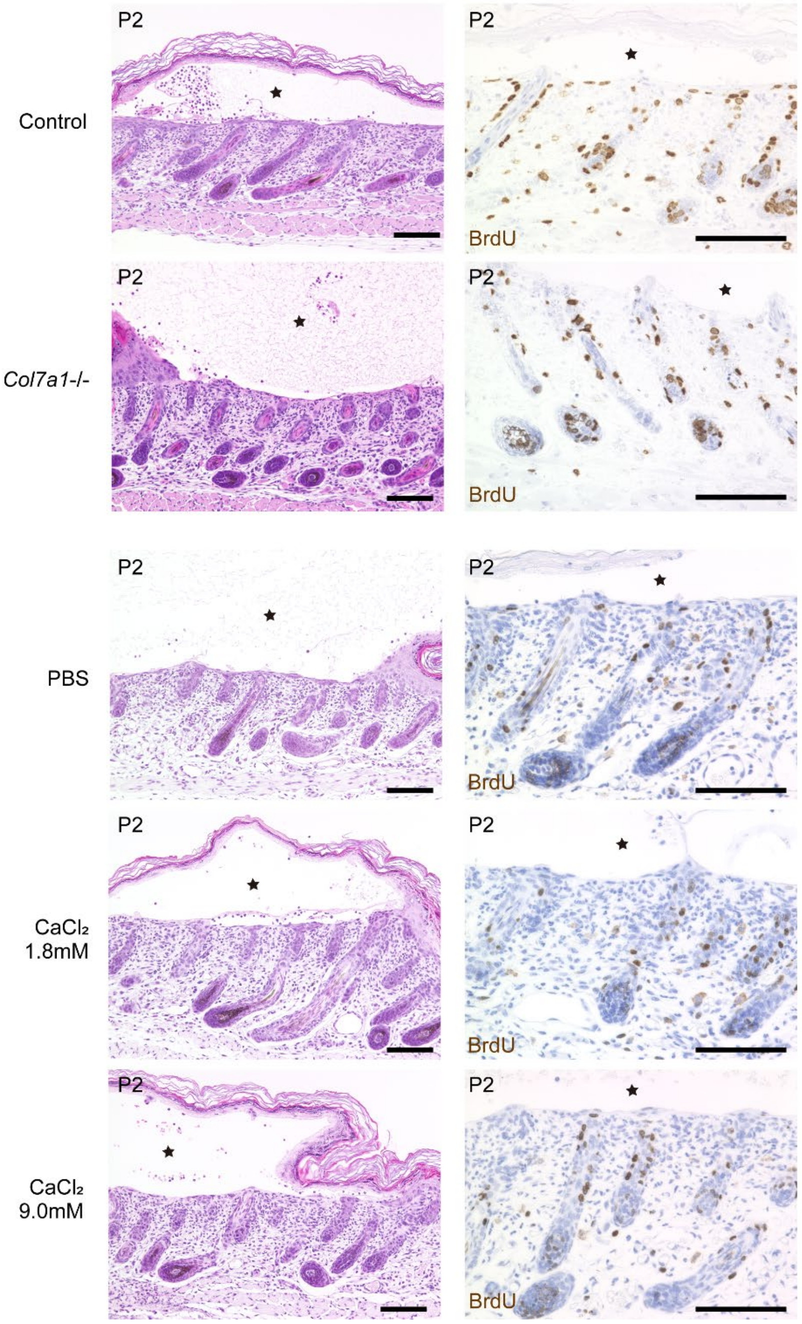


**Supplementary Figure 3. H&E and BrdU labeling of *Col7a1*-/- and Ca-treated blisters.**

H&E and BrdU labeling of blister samples at P2. Scale bar: 100 μm. Blisters are indicated by stars.

Supplementary Figure 4. Subepidermal blister healing in humans

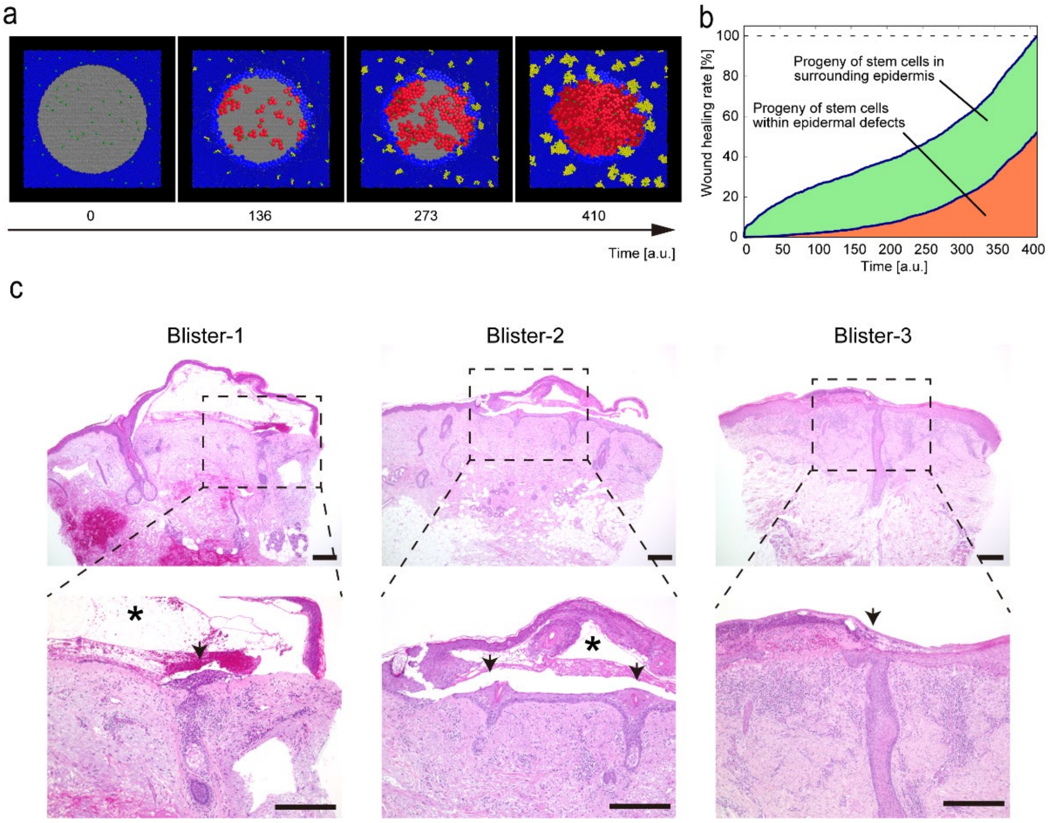


**Supplementary Figure 4. Subepidermal blister healing in humans.**

(a) A model of subepidermal blister healing in humans. SCs (green) give rise to progeny (simulating HF-derived cells; red) within epidermal defects or in the surrounding epidermis (IFE-derived cells; yellow). t: arbitrary time. See **Supplementary Video 4**. (b) Contribution of each progeny cell within the epidermal defect or of the surrounding epidermis to subepidermal blister healing in humans. (c) H&E staining of human subepidermal blister samples with re-epithelized areas (blisters 1, 2, and 3). Blisters are indicated by stars. The regenerated epidermis from HFs is indicated by arrows. Scale bar: 300 μm.

**Supplementary Table 1. Comparison of human and mouse HFs.**

| | Human (adult, body) | Mouse (P1, dorsal skin) |
|---|---|---|
| HF density | 14-32/cm$^2$ (1) | 12-19/1.2 mm epidermis (2) |
| HF diameter | 80-170 $\mu$m (1) | 40 $\mu$m (3) |
| HF depth | 1-4 mm (4, 5) | 100 $\mu$m (2) |

References to supplementary Table 1.

**Legends for Supplementary Videos**

**Supplementary Video 1. Mathematical modeling of subepidermal blister healing at the basal layer level (see Figure 6a).**

**Supplementary Video 2. Depletion of stem cells within epidermal defects in the mathematical model (see Figure 6c).**

**Supplementary Video 3. Effects of the impaired flattening of keratinocytes during epidermal regeneration in the mathematical model (see Figure 6e).**

**Supplementary Video 4. Mathematical modeling of subepidermal blister healing in humans (see Supplementary Figure 4).**
